# Heterotrophic bacteria drive sulfide oxidation in coastal sediments

**DOI:** 10.1101/2022.12.07.519552

**Authors:** Qun Cao, Yunyun Yang, Chuanjuan Lu, Qingda Wang, Yongzhen Xia, Qilong Qin, Luying Xun, Huaiwei Liu

## Abstract

Sulfate reduction and sulfur oxidation are very active in coastal sediments. They shape the biogeochemistry and microbial ecology at hot places of organic matter metabolism. Different from the well-studied sulfate reduction, sulfur oxidation in coastal sediments is still full of questions. Herein, we investigated the distribution of reduced sulfur compounds in differently layers of coastal sediments at the Yellow sea and found that sulfide (H_2_S), sulfane sulfur (S^0^), and thiosulfate mainly accumulated in anaerobic sediments and were mostly oxidized in anoxic and oxic interface in the sediments and the sea water. Bacterial community analysis indicated that heterotrophic bacteria are dominating species in surface sediments and sea water. Metagenome analysis showed that two sulfur-oxidizing genes encoding sulfide:quinone oxidoreductases (SQR) and persufide dioxygenases (PDO), were sharply more abundant than other sulfur-oxidizing genes in the coastal sediments. Since members of the marine Roseobacter clade were dominant in coastal waters and sediments, we studied the sulfur oxidation pathway in the Roseobacter *Ruegeria pomeroyi* DSS-3 and found that sulfide:quinone oxidoreductase, persulfide dioxygenase, and sulfite-oxidizing enzyme were the main enzymes for the oxidation of H_2_S, zerovalent sulfur, and sulfite/thiosulfate. This study, for the first time, clarified the dominating function of heterotrophic bacteria in sulfur oxidation in the coastal sediments and sea water.

**IMPORTANCE:** Coastal sediments are the most productive ecosystems. We performed the microbial community diversity and metagenomic analysis of seawater and coastal sediments of the Yellow Sea and explored the sulfur oxidation process in them. We found that heterotrophic bacteria are dominating species in surface sediments and sea water, sulfide and sulfane sulfur were mostly oxidized in surface sediments, and the genes encoding SQR, PDO, and SOE are abundant. Using *Ruegeria pomeroyi* DSS-3 as the model strain, we studied how these enzymes cooperate to oxidize H_2_S to sulfate. Thus, this research revealed the critical role of heterotrophic bacteria in sulfur oxidation in coastal sediments and sea water.

## INTRODUCTION

Coastal sediments are the most active places for sulfate reduction and sulfide oxidation. The continental shelf (0–200 m water depth) comprises only 8% of the global ocean area; however, 70%~80% of the global marine sulfate reduction takes place on it (1, 2). The shelf is very active for organic matter mineralization, and about half of the mineralization is driven by sulfate reduction (3). Hydrogen sulfide (H_2_S) is the end product of sulfate reduction, about 11.3 teramoles of sulfate are reduced to H_2_S every year (4). In oxygen rich surface, the produced H_2_S is re-oxidized to sulfate. Therefore, sulfate reduction and sulfur oxidation constitute a sulfur cycle, which is a major determinant of the biogeochemistry and microbial ecology in sulfate-rich coastal sediments (5, 6).

Microorganisms are the dominating driver of sulfur cycle in coastal sediments. In most coastal sediments, oxygen is depleted within the surface centimeters (7). Sulfate reduction mostly happens in anaerobic deep sediments, which is driven by sulfate reducing microorganisms (SRM) (8, 9). Most SRM are anaerobic bacteria with catabolic capacities for a wide spectrum of fermentation products including H_2_ and volatile fatty acids (VFAs) (10). Dissimilatory sulfite reductase (Dsr) is the functional marker for SRM. This enzyme catalyzes the reduction of sulfite to H_2_S (11). Different from SRM, most sulfur-oxidizing microorganisms (SOM) are aerobic bacteria residing in surface sediments. They are capable of catalyzing oxygen-dependent sulfide oxidation at rates that are orders of magnitude higher than the iron-driven chemical oxidations (12, 13). Some special SOM, including autotrophic/mixotrophic *Thiobacillus, Beggiatoaceae*, and cable bacteria, get focused attention due to either their CO_2_-fixing ability, conspicuous morphologies, or fascinating lifestyles (14–17). However, these bacteria occur in high abundances only in certain habitats; whereas 16S rRNA data suggest that SOM in most costal sediments are far exceed the currently known diversity.

Our previous studies indicate that heterotrophic bacteria with H_2_S-oxidizing capability are widely present in various habitats, including farms, forests, lakes, and coastal waters (18). The sulfur oxidation pathways they harboring are quite complex, but roughly can be divided into three segments: H_2_S oxidation to sulfane sulfur (S^0^), S^0^ oxidation to sulfite/thiosulfate, and sulfite/thiosulfate oxidation to sulfate (19, 20). For the first segment, sulfide-quinone oxidoreductase (SQR) and flavocytochrome c sulfide dehydrogenase (FCC) are central enzymes. For the second segment, persulfide dioxygenase (PDO) is the main enzyme. Sulfur oxidation system (SOX) and sulfite-oxidizing enzyme (SOE) are responsible for the last segment. Except for these unambiguous pathways, SOX catalyzing S^0^ oxidation directly to sulfate (21), thiosulfate dehydrogenase A (TsdA) catalyzing thiosulfate oxidation to tetrathionate (22), and APS reductase (APR) catalyzing sulfite oxidation to sulfate were also reported (23), but their ecological roles need further investigation. Most SOM contain one or two segments of the sulfur oxidation pathway, but bacteria harboring an integrated pathway are also widely present.

Herein, we investigated the sulfur oxidation process, the participating SOM, and the functioning genes in coastal sediments of the Yellow Sea. The sampling location (N36.373534, E120.70228) is in Qingdao, China (Fig. 1A), and the time was in mid-August, a period when *Enteromorpha prolifera* outbreaks and brings abundant organic matters to this coastal zone every year. We found that sulfide and sulfane sulfur were mostly oxidized in surface sediments (0~10 cm). As in terrestrial environments, heterotrophic bacteria dominated the oxidation process. Metagenomic analysis indicated that genes coding for SQR, PDO, and SOE are abundant. To elucidate how these enzymes are working together for H_2_S oxidation, we used a rhodobacterium that harbors these genes as a model and studied the pathway of sulfide oxidation to sulfate in it.

**Fig. 1.**
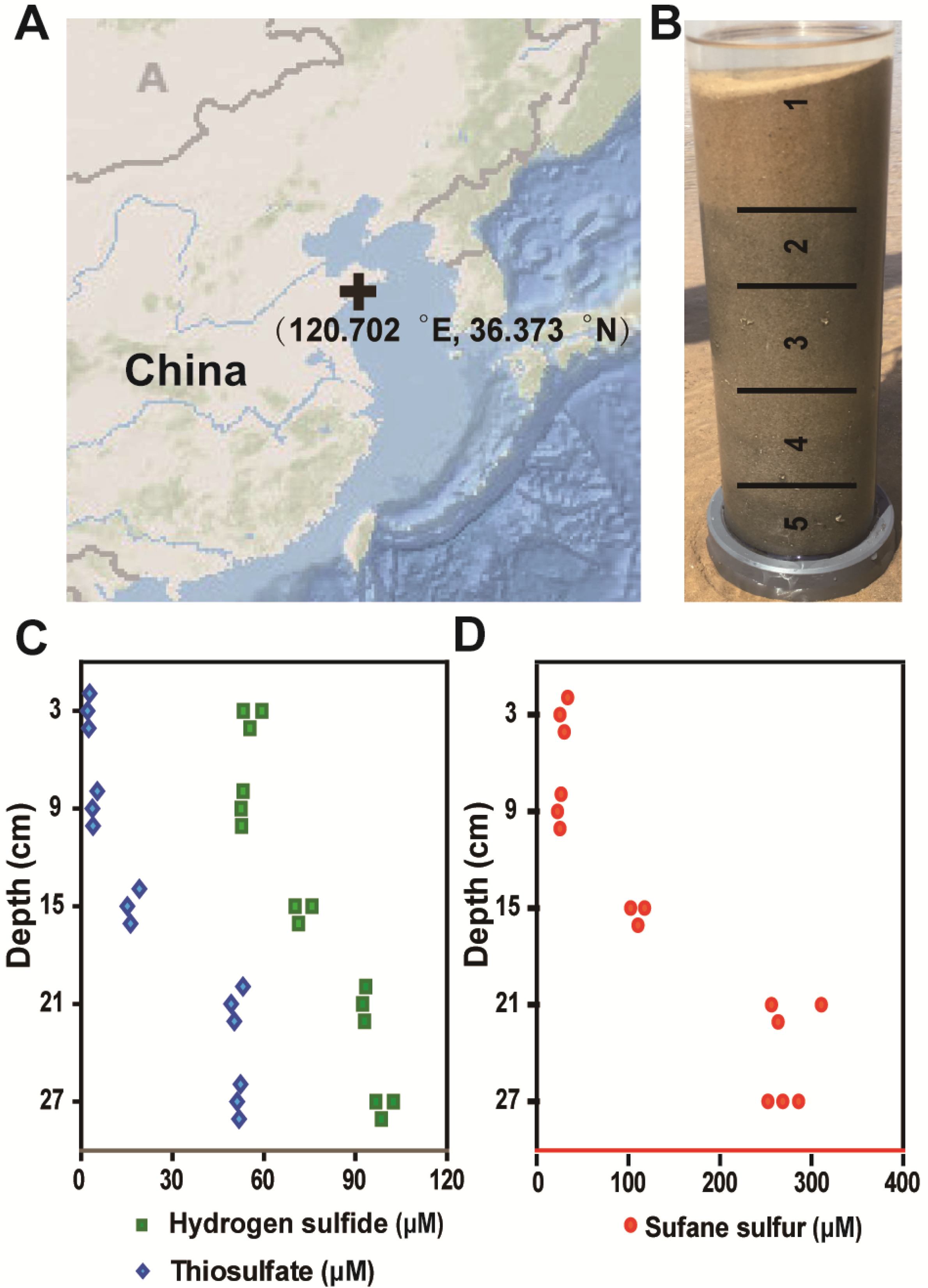
Analysis of sulfur containing compounds in coastal sediments. (A) Geographic location of the sampling station. (B) Sediment column was divided into five segments according to their color. (C, D) Sulfur containing compounds in each sediment were quantified. The ordinate is the average depth of analyzed sediments. Three parallel samples were collected and analyzed (n = 3).

## RESULTS

### Sulfide and sulfane sulfur were richer in deep sediments than in surface ones

To maintain the sampled sediments undisturbed, we inserted a glass tube (7.5 cm in diameter) into sediments deep to 30 cm to take out a sediment core. The sediment core was stratified in color—deeper layer was obviously darker than surface one (Fig. 1B). We divided the sediment column into five segments according to their colors and took three samples from each segment to quantify contents of sulfane sulfur, H_2_S, and thiosulfate. Samples from surface segment (3~5 cm in depth) contained 29.7 μM sulfane sulfur, 55.9 μM H_2_S, and 2.4 μM thiosulfate. In contrast, samples from the deeper segment (27 cm in depth) contained 268.8 μM sulfane sulfur, 99.2 μM H_2_S, and 51.8 μM thiosulfate. Sulfane sulfur and H_2_S were not detected in the sea water above the sediment core. Thiosulfate content in the sea water was similar as in the surface sediments. Therefore, the oxidation of thiosulfate, sulfane sulfur, and H_2_S in the surface sediment from the deep sediment was 96%, 89%, and 44%, respectively. The residual sulfane sulfur and H_2_S were likely oxidized in the interface of sea water and sediment, as they were undetectable in the surface water.

Bio-generated sulfane sulfur exists in many forms including HS_n_H and RS_n_H (n≥2), RS_n_R (n≥3), and S_8_. S_8_ was the dominating species in both surface and deep sediments (Fig. S1). Sulfite was not detected in the sediments or sea water.

### Heterotrophic bacteria were abundant in both surface sediments and above sea water

To investigate the functional sulfur-oxidizing strains in surface sediments and sea water, we analyzed the bacterial community in them (methods are provided in supplementary information). The bacterial species showed large diversity in both sediments and sea water. In details, *Bacillales, Flavobacterials*, *Nevskiales, Rhodobacteriales*, and *Candidatus* Actinomarinales were top 5 most abundant orders in surface sediments. Together, they counted for 57.5% of the total bacterial community (Fig. 2A). Differently, *Rhodobacteriales*, *Flavobacterials, Pseudomonadales, Candidatus Actinomarinales*, and SAR116 cluster were top 5 most abundant orders in sea water (Fig. 2B). Together, they counted for 65.5% of the total bacterial community. The data also indicated that heterotrophic bacteria were dominating strains in both the surface sediments and sea water. They counted for 91.9 % of the total bacteria in the surface sediments and 95.5% in the sea water.

**Fig. 2.**
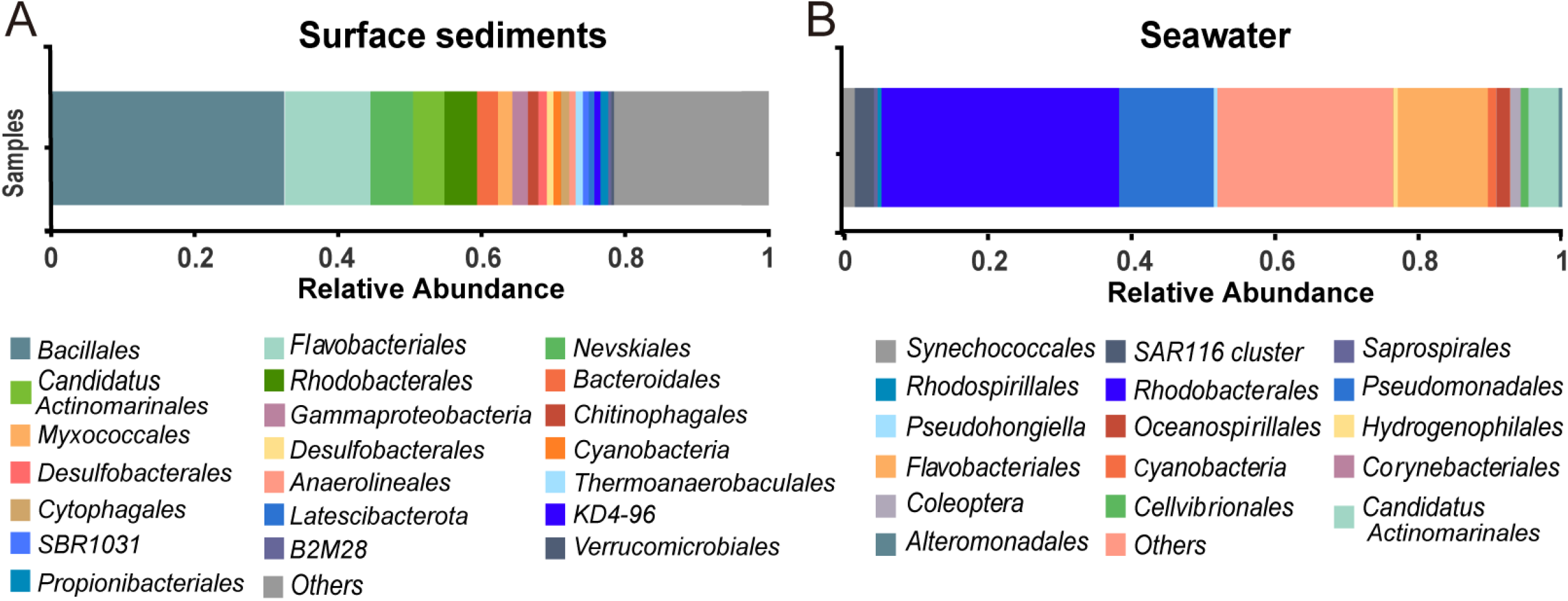
Bacterial community in surface sediments (A) and above sea water (B).

### Sulfur metalizing genes were rich in the surface sediments

We performed metagenomic sequencing to analyze genomes in surface sediments (methods are provided in supplementary information). Our data showed that metabolism genes were overwhelming abundant, counting for 79.6% of the total genes (Fig. 3A). Of the top 20 most abundant biogeochemical cycles related genes, 17 of them were sulfur metabolism related (Fig. 3B, except for *napA, narG*, and *nosZ*. The three are nitrogen metabolism related). The sulfur oxidizing gene *pdo* and *sqr* were ranked as 6^th^ and 7^th^ among all genes. To get a clear picture, we selected all sulfur metabolism related genes out and divided them into three categories: sulfur oxidation, sulfur reduction, and organic sulfur metabolism. For the sulfur oxidation category, *sqr* is more abundant than *fcc* genes *fccA* and *fccB*), suggesting that SQR is the main H_2_S oxidizing enzyme in surface sediment (Fig. 4A). The *pdo* gene is almost as abundant as *sqr*, and much more abundant than *sox* genes, implying that PDO is the main zero-valent sulfur oxidizing enzyme. The *soe* genes are as abundant as the *sox* genes. The sulfur oxygenase reductase encoding gene *sor* (found in chemolithoautotrophic crenarchaeote) and the thiosulfate dehydrogenase encoding gene *tsdA* (found in purple sulfur bacterium) are in much lower abundance compared to others, indicating that they are not the main players in the sulfur oxidation process in the coastal region.

For the sulfur reduction category, the sulfate permease gene *sulP* is the most abundant (also the top 1 abundant gene in whole metagenomic data) (Fig. 4B). Sulfate reducing genes *sat* and *aprA* are the second and third abundant, respectively. Sulfite reductase genes including *aprA, dsrA*, and *dsrB* are little lower in abundance. These data suggest that sulfate reduction process is also active in the surface sediments. For the organic sulfur metabolism, cysteine synthase gene *cysK* and methionine synthesis related gene *metY* are top 2 most abundant (Fig. 4C).

**Fig. 3.**
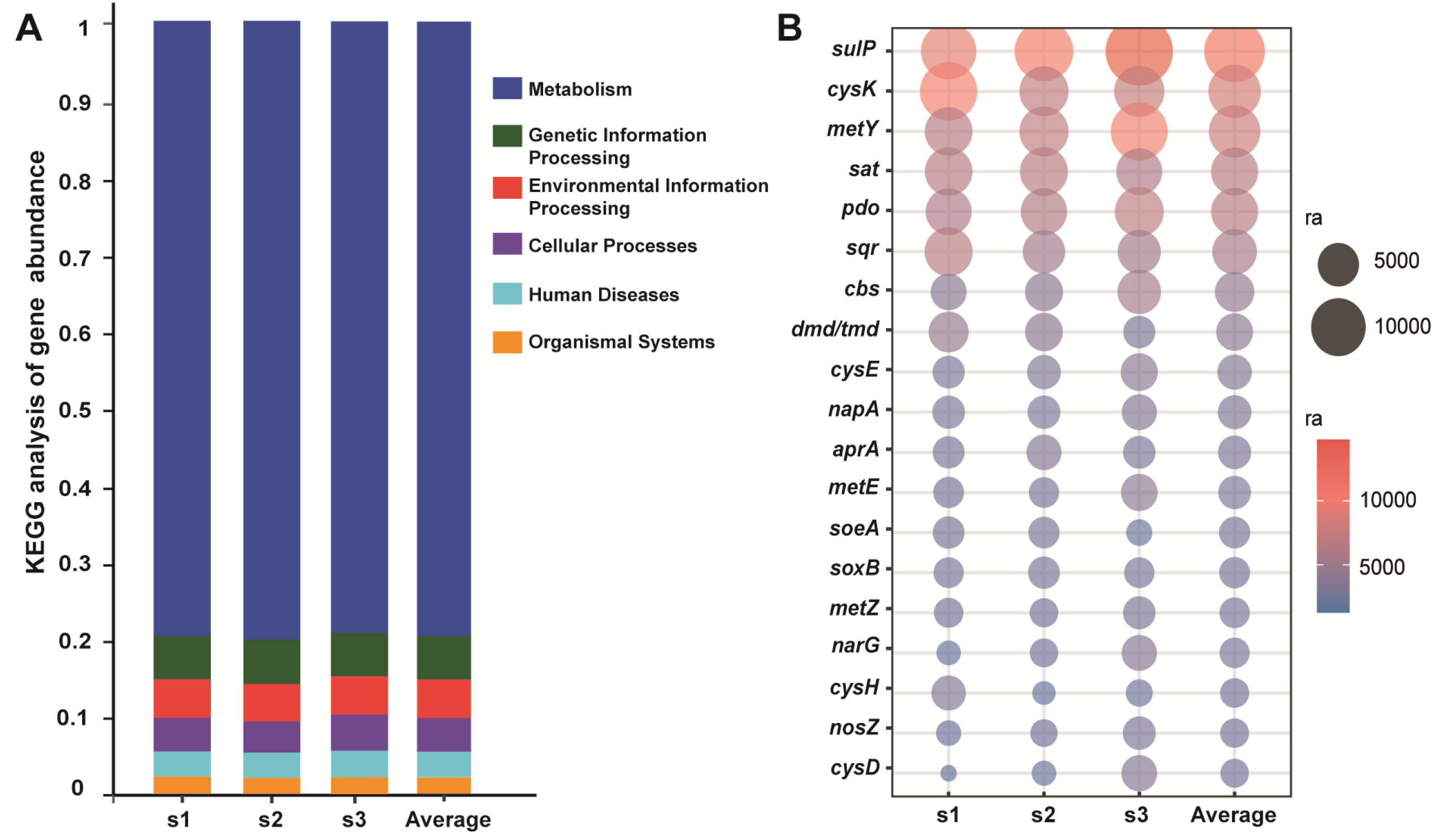
Metagenomic analysis of surface sediment samples. (A) KEGG analysis showed that metabolic genes were the most abundant category. (B) Top 20 most abundant genes related to biogeochemical cycles. 17 of 20 are sulfur metabolism related. Three parallel samples (s1~s3) were collected and analyzed (n = 3). Average data of these three samples were shown.

**Figure 4.**
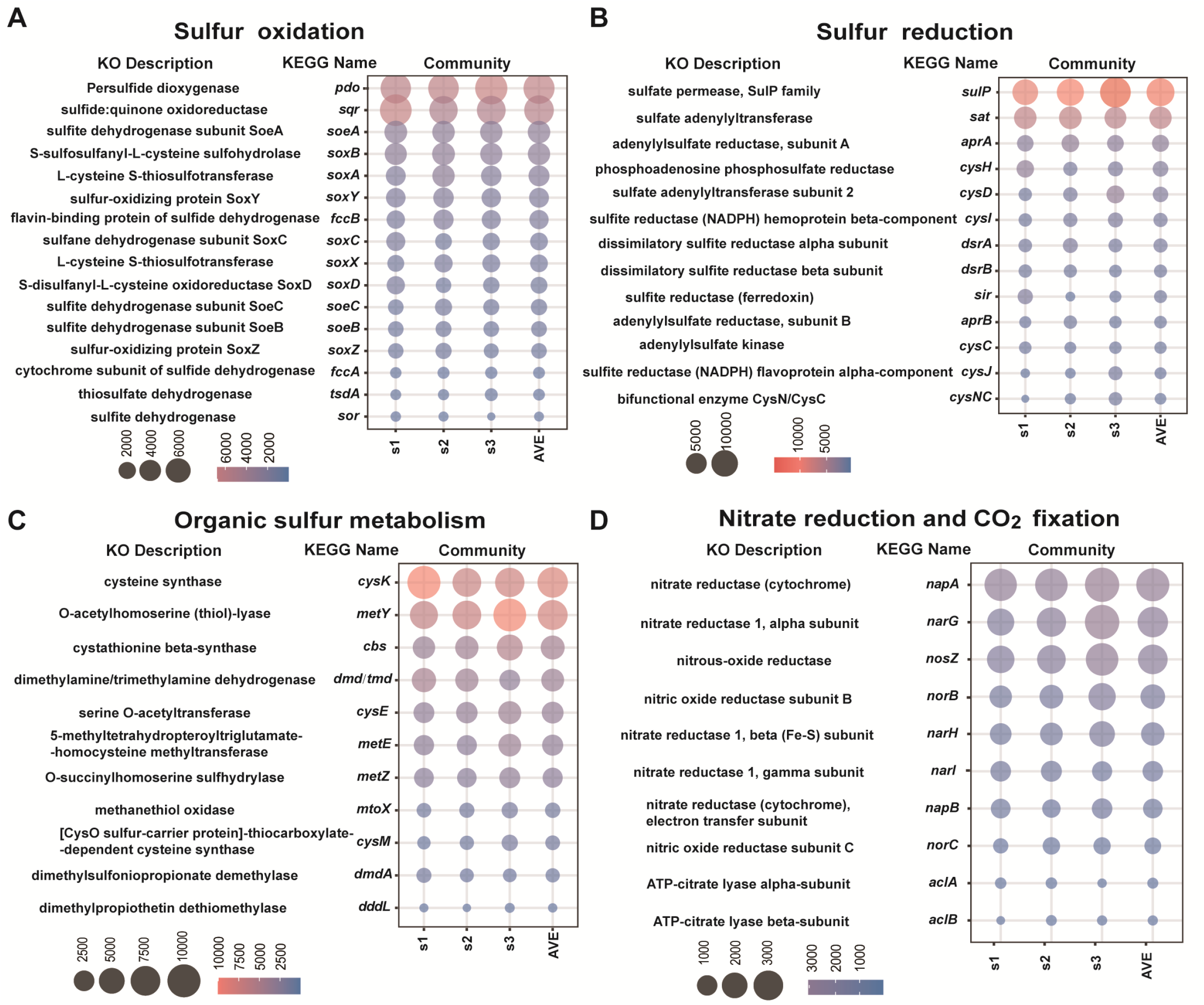
Abundance analysis of sulfur oxidation (A), sulfur reduction (B), organic sulfur metabolism (C), and nitrate reduction and CO_2_ fixation (D) genes in surface sediments. Three parallel samples (s1~s3) were collected and analyzed (n = 3). Average data of these three samples were shown.

Dimethylsulfoniopropionate (DMSP) degrading-enzyme genes (*dmd/tmd, dmdA*, and *mtoX*) are also present but in lower abundance than that of the inorganic sulfur metabolizing genes.

In addition to sulfur metabolism genes, we also listed nitrate reduction and CO_2_ fixation related genes out (Fig. 4D). The nitrate reduction related genes are present in modest abundance and CO_2_ fixation related genes (*aclA and aclB*) are in low abundance. Taken together, the metagenomics data indicate that sulfur metabolism genes from heterotrophic bacteria are richer than those from autotrophic bacteria, and the sulfur oxidation coupled CO_2_ fixation process (driven by autotrophic bacteria) may rarely happen in surface sediments.

### Distribution of the sulfur oxidation enzymes in rhodobacteriales

Since *Rhodobacteriales* are rich in both surface sediments and sea water, we wondered whether they have sulfur oxidation capability (methods are provided in supplementary information). From 1904 sequenced *Rhodobacterales* genomes (NCBI database, updated to 6 June 2022), we found 344 SQR encoding genes in 289 *Rhodobacterales* and 284 FccB encoding genes in 260 *Rhodobacterales* (Fig. 5A). SQR and PDO coverages in *rhodobacteriales* were 15.2 % and 13.7%, respectively (Table S4). Most of SQRs belong to type II and most of FccBs belong to subgroup 1 (Fig. S2 and S3). Compared to H_2_S oxidation enzymes, sulfane sulfur oxidation enzyme PDO is more widely present in *Rhodobacterales*, evidenced by 836 PDO from 592 *Rhodobacterales* with a coverage of 31.1% (Fig. 5B). Most of PDO belong to the type II (Fig. S4). SOE and SOX encoding genes were also observed, with coverages range from 12.4% to 21.4% (Table S4).

**Fig. 5.**
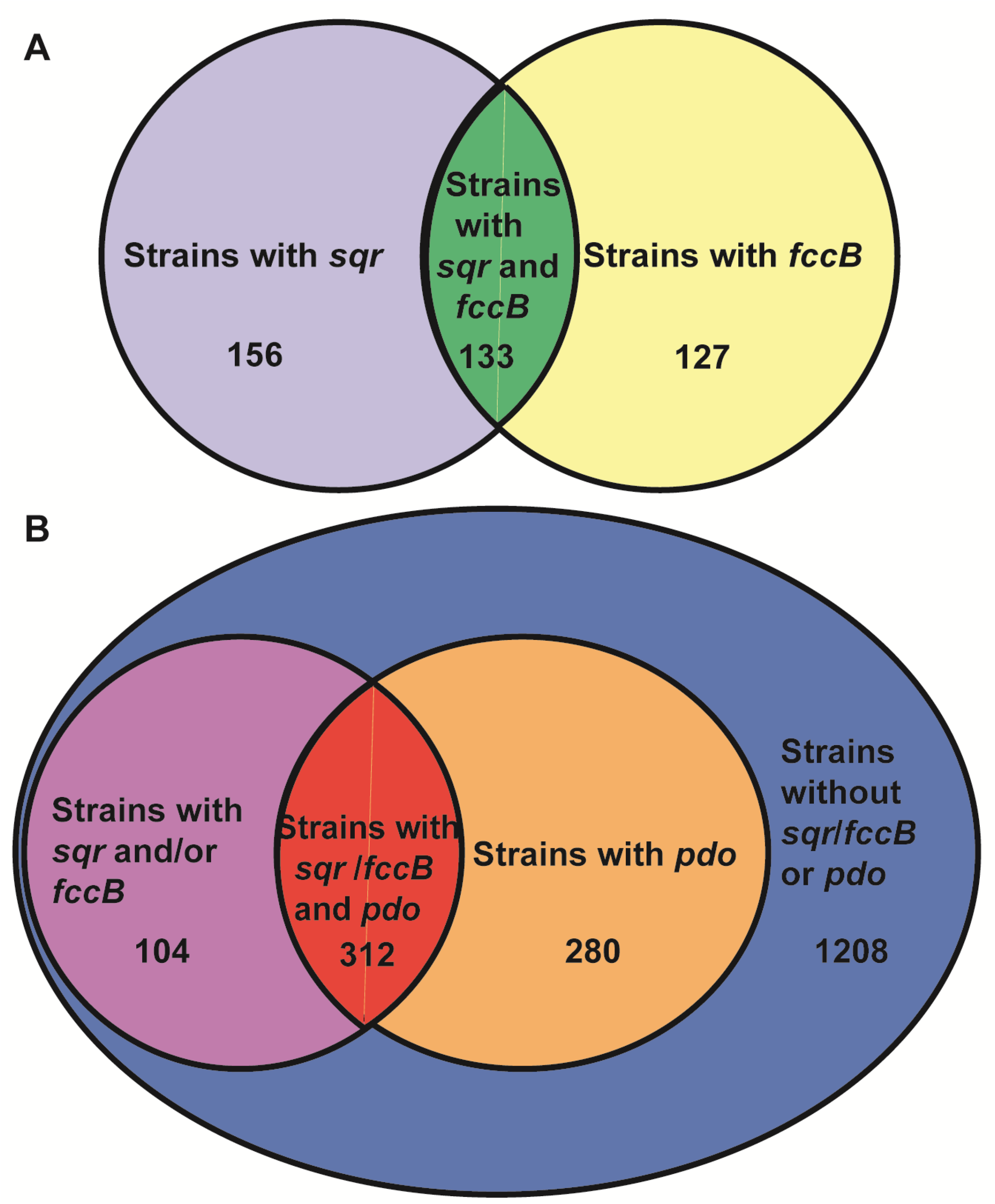
The distribution of *sqr, fccB* (A), and *pdo* (B) genes in sequenced 1904 Rhodobacterales genomes. 31.1% Rhodobacterales have *pdo*, 15.2% have *sqr*, 13.7% have *fccB*. 6.9% have both *sqr* and *fccB*. 16.4% have *sqr/fccB* and *pdo*.

### Characterizing the sulfur oxidation pathway in *R. pomeroyi* DSS-3

To investigate how the sulfur oxidizing enzymes cooperate in H_2_S/sulfane sulfur oxidation. We studied a model bacterium of *Rhodobacterales, R. pomeroyi* DSS-3. Genome analysis indicated that this strain has an integrated sulfur oxidation pathway composed by SQR, Fcc, PDO, SOX, and SOE (Fig. 6A).

**Fig. 6.**
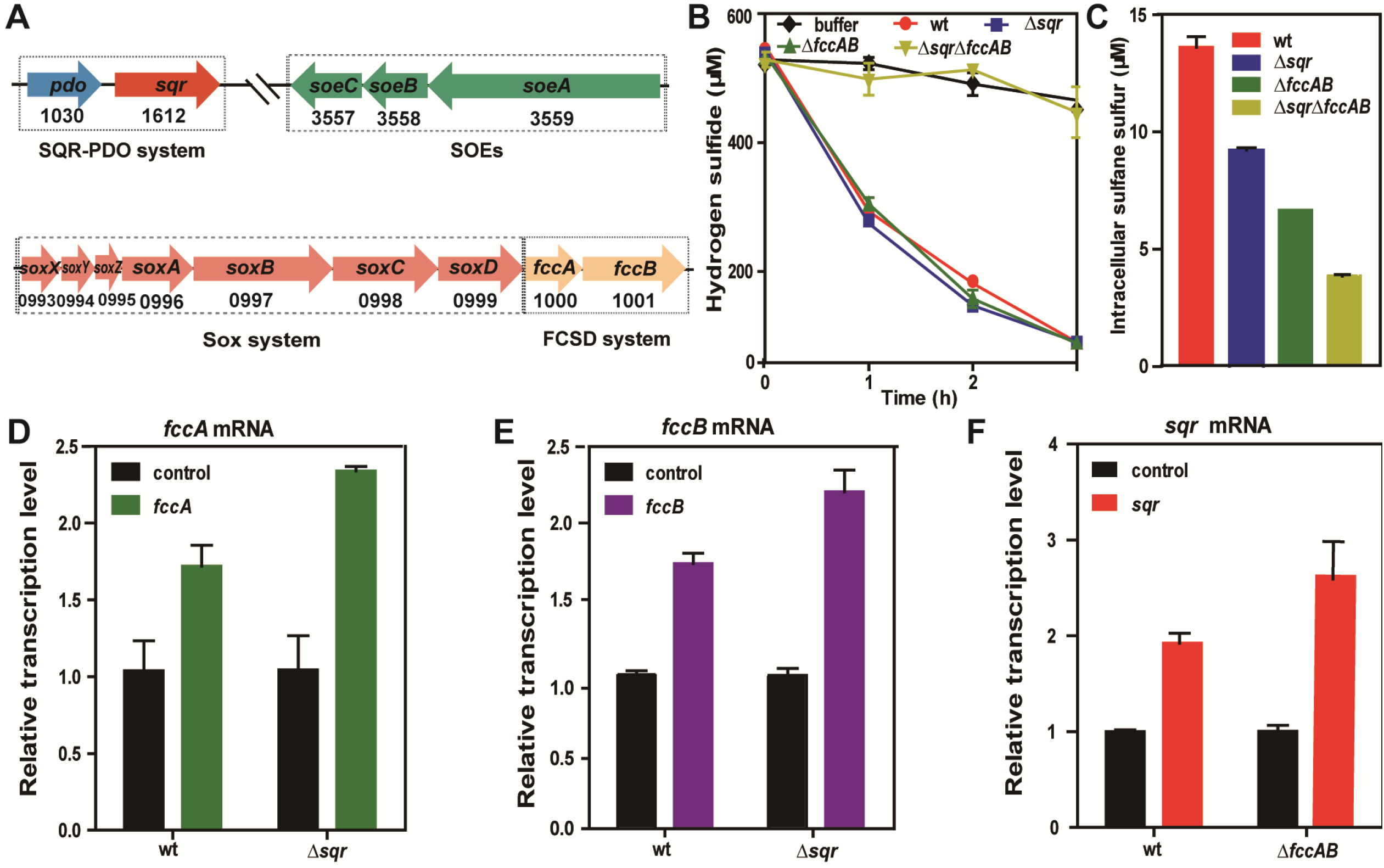
Sulfur oxidizing genes in *R. pomeroyi* DSS-3. (A) Locations of the sulfur-oxidizing genes in *R. pomeroyi* DSS-3 genome. The complete *R. pomeroyi* DSS-3 genome is from GenBank database with accession number NC_003911.12. (B) *sqr* or *fccAB* single deletion strains showed no impaired activity on H_2_S oxidation while double deletion strains totally lost the activity. (C) Sulfane sulfur production from H_2_S was obviously lower in double deletions strain than that in wt and single deletion strain. (D,E,F) Analysis of *fccA, fccB* and *sqr* transcription with qRT-PCR. Transcription of *fccA* and *fccB* was higher in *Δsqr* than in wt and transcription of *sqr* was higher in *ΔfccAB* than in wt. Data were from three independent experiments, shown as average with s.t.

First, we constructed three mutant strains of *R. pomeroyi* DSS-3: *Δsqr, ΔfccAB*, and *ΔsqrΔfccAB*. We tested their H_2_S oxidation rates and observed that compared to *R. pomeroyi* DSS-3 wild type strain (wt), *Δsqr* and *ΔfccAB* showed no apparent difference on H_2_S oxidation (Fig. 6B). Whereas, *ΔsqrΔfccAB* almost lost the ability of H_2_S oxidation. Since both SQR and FCC oxidize H_2_S to sulfane sulfur, we analyzed the intracellular sulfane sulfur during H_2_S oxidation process. *ΔsqrΔfccAB* accumulated apparently lower intracellular sulfane sulfur than wt, *Δsqr*, and *ΔfccAB* (Fig. 6C). These results indicated that both SQR and *FccAB* are involved in H_2_S oxidation, and these two are functionally complementary.

To test whether expression of *sqr* and *fccAB* is inducible by H_2_S, we analyzed their mRNA level using RT-qPCR. We found that mRNA levels of *sqr* and *fccAB* were both increased after H_2_S induction (Fig. 6D~6F). Further, in *ΔfccAB*, the *sqr* mRNA level increased more than that in wt. Likewise, in *Δsqr*, the *fccAB* mRNA level increased more than that in wt. These results demonstrated both *sqr* and *fccAB* are inducible. Once any one is absent, the other will increase its expression.

Second, we constructed a PDO deletion strain *Δpdo* and tested its ability on hydrogen persulfide (HSSH, sulfane sulfur containing compound) oxidation. Compared to wt, *Δpdo* lost more than 50% ability of HSSH oxidation (Fig. 7A) The main oxidation product was thiosulfate (Fig. 7B). Sulfite production was also detected, but at much low levels (Fig. 7C). During the first 3 hours, no obvious sulfate production was detected. After 3 hours sulfate production sharply increased, indicating that there was a lagging time for thiosulfate/sulfite being further oxidized to sulfate (Fig. 7D). Interestingly, *Δpdo* also showed impaired ability on H_2_S oxidation (Fig. 7E), but no obvious change on thiosulfate (Fig. 7F) and sulfite oxidation (data not shown). Third, a SOE deletion mutant *ΔsoeABC* and a SOX deletion mutant *ΔsoxYZ* were constructed. The *ΔsoeABC* mutant showed a little impaired activity on sulfite oxidation (Fig. 7G), but severely impaired activity on thiosulfate oxidation (Fig. 7H). Compared to wt, *ΔsoxYZ* showed no obviously difference on H_2_S, HSSH, or sulfite oxidation (data not shown). Its thiosulfate oxidation activity was a little impaired (Fig. 7I). These results indicated that SOE has both thiosulfate and sulfite oxidation activity with the former is its main function. SOX system cannot oxidize sulfite and its thiosulfate oxidation activity is limited compared to SOE.

**Fig. 7.**
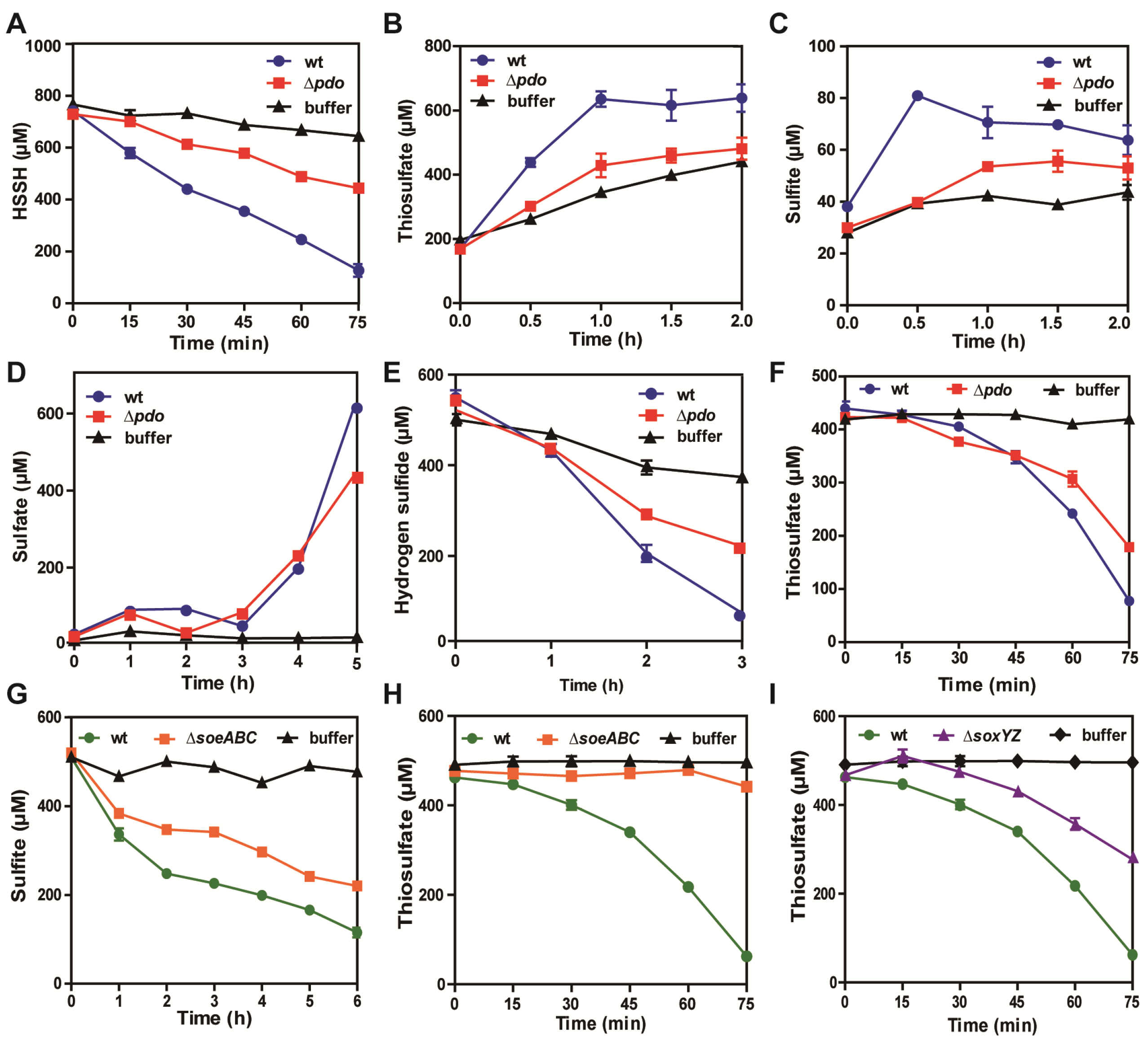
Sulfane sulfur, sulfite, and thiosulfate oxidation activity assay of *R. pomeroyi* DSS-3. (A) *pdo* deletion severely impaired HSSH oxidation. (B) Thiosulfate production from HSSH oxidation. (C) Sulfite production from HSSH oxidation. (D) Sulfate production from HSSH oxidation. (E) *pdo* deletion also impaired H_2_S oxidation. (F) *pdo* deletion had no obvious influence on thiosulfate oxidation. (G) *soe* deletion impaired sulfite oxidation. (H) *soe* deletion severely impaired thiosulfate oxidation. (I) *sox* deletion impaired thiosulfate oxidation.

## DISCUSSION

In anoxic deep sediments of the ocean, anaerobic microorganisms use sulfate as an electron acceptor to drive the organic carbon degradation. In oxygen rich surface sediments and sea water, aerobic microorganisms oxidize H_2_S/sulfane sulfur back to sulfate (Fig. 8A). Without the sulfur oxidation process, sulfate reduction cannot last and finally organic carbon degradation will be impeded. Therefore, sulfur oxidation is equally important for the oceanic carbon cycle as sulfate reduction. In this study, we investigated the sulfur oxidation process in coastal sediments of the Yellow sea. We found that sulfane sulfur, H_2_S, and thiosulfate were rich in deep sediments but not in surface ones. Bacterial community analysis indicated that heterotrophic bacteria were dominating species in surface sediments and sea water. Metagenome analysis revealed that sulfur oxidizing enzymes including SQR, PDO, and SOE were abundant in surface sediments. Using *R. pomeroyi* DSS-3 as a model bacteria, we found that these enzymes were responsible for sulfane sulfur/H_2_S oxidation to sulfate (Fig. 8B). The autotrophic/mixotrophic *Thiobacillus* were rare in surface sediments (0.13%). No cable bacteria were present in them either. Although *Gammaproteobacteria* were present in considerable abundance (13.5%), CO_2_ fixation related genes *aclA and aclB* were very low, indicating that the sulfur oxidation coupled CO_2_ fixation process driven by *Gammaproteobacteria* rarely happened in surface sediments. Considering the dominating abundance of heterotrophic bacteria in both surface sediments and sea water, and the overwhelming richness of sulfur oxidizing genes, we believe that heterotrophic bacteria drive the sulfur oxidation process in the surface sediments and above sea water, as they do in many terrestrial environments.

**Fig. 8.**
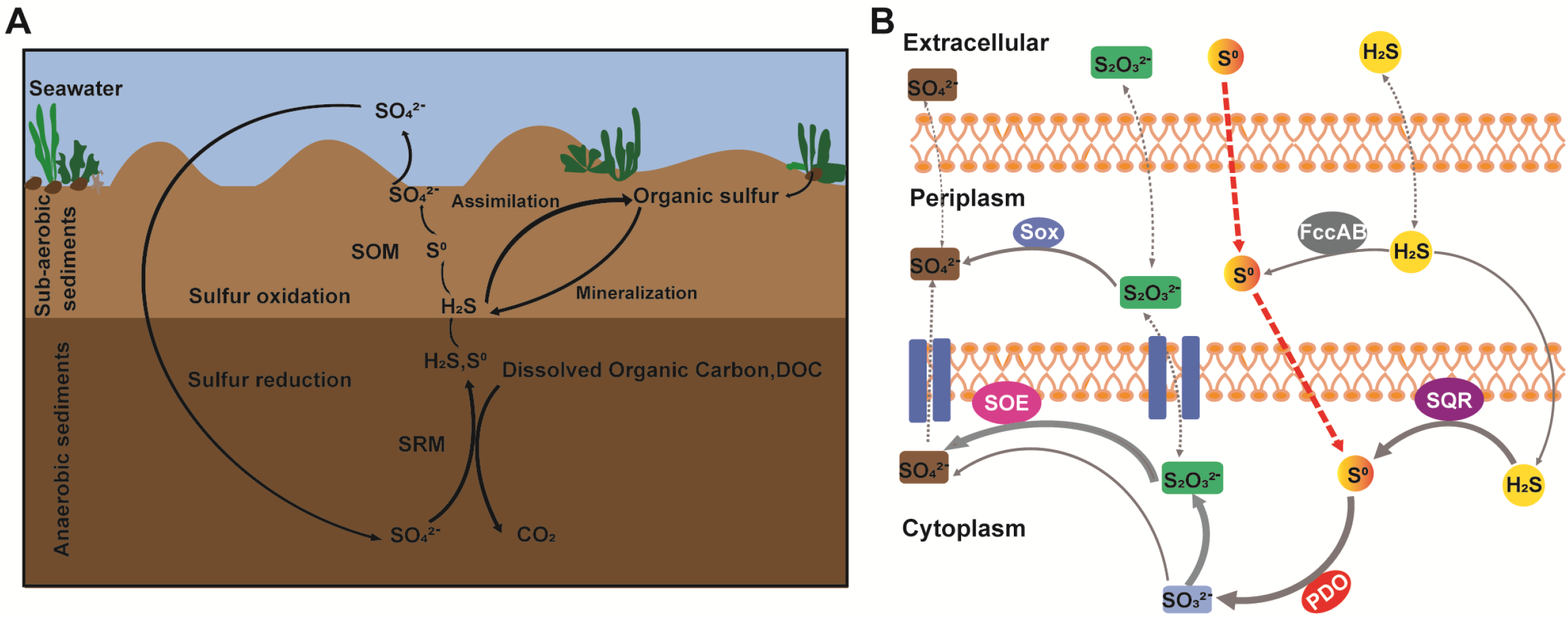
Sulfur cycling in coastal sediments (A) and the main sulfur oxidation pathway. (A) The reduction of sulfate to sulfide and sulfane sulfur by SRM is accompanied by mineralization of organic carbon in anaerobic sediments. Reduced sulfide and sulfane sulfur get oxidized in surface sediments and sea water by SOM. (B) H_2_S is oxidized to sulfane sulfur by SQR and FccAB, SQR plays a major role in this step; PDO oxidizes sulfane sulfur to sulfite, which spontaneously reacts with sulfane sulfur to generate thiosulfate. SOE oxidizes thiosulfate and sulfite to sulfate directly. A small portion of thiosulfate is transported to the periplasmic space and gets oxidized by Sox system.

Sulfane sulfur is more abundant than H_2_S in coastal sediments. This phenomenon may be due to two reasons. First, H_2_S is oxidized faster than sulfane sulfur; second, sulfane sulfur is the main product of sulfate reduction. Previous studies indicated that *K_m_* values of PDO are much higher than those of SQR (24, 25), and microorganisms commonly contain more sulfane sulfur than H_2_S inside cells (26, 27). These make the first reason highly possible. However, it cannot explain the richness of sulfane sulfur in anaerobic deep sediments. On the other hand, our recent study indicated that for fission yeast cysteine synthase, sulfane sulfur is a better substrate than H_2_S (28). Blocking sulfane sulfur supply caused cysteine synthesis deficient at the presence of abundant H_2_S. Therefore, the classic pathway of sulfate reduction to H_2_S, and then H_2_S is used for cysteine synthesis is challenged. It is possible that sulfate is reduced to sulfane sulfur, and the latter is easily reduced to H_2_S. Thus, H_2_S is efflux of sulfane sulfur. If the assimilatory sulfate reduction works this way, dissimilatory sulfate reduction produces sulfane sulfur is reasonable. Further investigations are required for judging this assumption.

*sqr* is much more abundant than *fccAB* in surface sediments, indicating that the former is a main player in H_2_S oxidation. The latter may function as a substitute. *pdo* is the richest sulfur oxidizing gene, following by *sqr*. Studies in *R. pomeroyi* DSS-3 demonstrated that PDO is the key enzyme. Its deletion led to both sulfane sulfur and H_2_S oxidation deficiency. We detected thiosulfate but not sulfite in sediments/sea water, and for *R. pomeroyi* DSS-3, thiosulfate is the main product of H_2_S and sulfane sulfur oxidation. These results indicated that sulfite can quickly react with sulfane sulfur to produce stable thiosulfate. *soe* genes are more abundant than *sox* genes in surface sediments and SOE is the main thiosulfate oxidizer in *R. pomeroyi* DSS-3. This finding is interesting because SOE commonly was deemed as a sulfite oxidase in previously (15, 29). We indeed found that SOE also has sulfite oxidation activity whereas SOX does not. Currently, there is no clearly discernible functional molecular marker for SOM, our study indicated that SQR, PDO, and SOE can be used as molecular markers for identifying marine SOM.

## MATERIALS AND METHODS

### Sample collection

Samples were collected from Aoshan Bathing Beach (N36.373534, E120.70228) of Qingdao, China. The sampler was a glass tube with a diameter of 7.5 cm. We inserted the sampler into beach sediments deep to 30 cm, and then took out the sediment without disturbing its layers. Since the sediment column taken out was stratified in color, we divided the column into five layers according to the color change. To quantify sulfur containing compounds in different layers, sediments were taken out from each layer and weighed. Tris-HCl buffer (50 mM, pH 9.5) was added with equal weight to dissolve compounds of sediments into the buffer.

### Quantification sulfur containing compounds

Total sulfane sulfur in sediments was quantified with a previously reported method (26). Briefly, 1 mM sodium sulfite (NaSO_3_) was added into the sediment-Tris-HCl buffer mixture, and then boiled the mixture at 95°C for 10 min. This step converted unstable sulfane sulfur into stable thiosulfate. After the boiling step, the mixture was centrifuged at 11000 ×g for 5 min to obtain the supernatant. Monobromobimane (mBBr) was added into the supernatant to derivatize thiosulfate (Na_2_S_2_O_3_). Quantification of the mBBr derived thiosulfate was performed via HPLC.

To quantify H_2_S, the supernatant was directly derivatized with mBBr and analyzed via HPLC. For quantification of total sulfane sulfur and H_2_S in sea water, the experiment protocols were the same.

Sulfite and thiosulfate in sediments and sea water were quantified using an ion chromatograph system (ICS; model ICS-1100; Dionex, USA) in the anion detection mode. The column was an Ion Pac AS19 column and its temperature was set to 30°C. The eluent automatic generation device (RFC-30) and ASRS_4 mm suppressor were used to generate gradient elution of 1 ml/min KOH. Under these conditions, the elution times of sulfite and thiosulfate were 7.3 min and 20.8 min, respectively. For *Ruegeria pomeroyi* DSS-3 culture samples, H_2_S quantification was performed with the methylene blue method (30) and total sulfane sulfur quantification was performed with the cyanide method (31). The quantification of sulfite, thiosulfate, and sulfate was performed with the ion chromatograph method as described above. Different from sulfite and thiosulfate, sulfate had an elution time of 7.8 min.

### Determination of sulfane sulfur species in sediments

To determine the distribution of sulfane sulfur species in sediments, a previously reported method was applied (32). Briefly, the sample was derivatized with methyl trifluoromethanesulfonate to convert H_2_S_n_ to dimethylpolysulfide. Elemental sulfur (S_8_) cannot be derivatized but can become soluble in methyl trifluoromethanesulfonate-methanol solution. Both dimethylpolysulfide and S_8_ were detected by using HPLC with a UV detector.

### Strains, culture conditions, and chemicals

*R. pomeroyi* DSS-3 and its mutant strains used in this study are listed in Table S1. A modified YTSS medium (1/2 YTSS) containing 4 g/L yeast extract, 2.5 g/L tryptone, 20 g/L sea salts was used for their cultivation. The cultivation temperature was 30°C with shaking at 200 rpm. *R. pomeroyi* DSS-3 gene deletion strains were constructed by using a previously reported method (33, 34). Briefly, to delete a target gene, upstream and downstream fragments of the target gene were amplified by PCR. These two fragments were ligated with the linearized plasmid pK18mobsacB to construct a deletion plasmid. The deletion plasmid was transformed into *E. coli* MFD pir and then transferred to *R. pomeroyi* DSS-3 by conjugation. The transformed *R. pomeroyi* colonies were confirmed through colony PCR and DNA sequencing. All primers used in the deletion process are listed in Table S2.

Hydrogen persulfide (HSSH) was prepared following a reported method [19]. NaHS Na_2_SO_3_, Na_2_S_2_O_3_, NaSO_4_, and other chemicals was purchased from Sigma-Aldrich (MO, USA).

### RNA extraction and qRT-PCR analysis

*R. pomeroyi* DSS-3 cells at log phase (OD_600_ reached 0.7~1.2) were harvested by centrifugation at 4000×g, 4°C, for 10 min. Total RNA was isolated by using the TaKaRa MiniBEST universal RNA extraction kit, and the concentration of RNA was determined with Nanodrop K5500 (Kaiao, Beijing, China). The cDNA was acquired by using the Prime Script RT reagent kit with genomic DNA (gDNA) eraser (TaKaRa, Beijing, China). The SYBR Premix Ex Taq II kit (TaKaRa) was used for qRT-PCR, and the reactions were run in a Light Cycler 480 II sequence detection system (Roche, Shanghai, China). The *rpoC* and *gyrA* genes were used as reference. Primers used for real-time PCR are listed in Table S3.

### Sulfur oxidation activity of *R. pomeroyi* DSS-3

The strain was cultured in 1/2 YTSS medium overnight. The overnight culture (1 ml) was transferred into 100 ml of fresh medium and cultured to OD_600_= 2.0. Cells were collected by centrifugation (4,000×*g*, 5 min) and resuspended in Tris-HCl buffer (pH 8.4, 50 mM) at OD_600_=5.0. NaHS (500 μM), HSSH (800 μM), Na_2_SO_3_ (500 μM), or Na_2_S_2_O_3_ (500 μM) was added into 10 ml cell suspension in a 50-ml scale tube. The tube was then sealed with a silicone cap and incubated at 30°C with shaking (100 rpm). The oxidation products were measured as mentioned above.

## Data Availability Statement

Bacterial 16S rDNA raw data have been deposited in NCBI Sequence Read Archive (SRA) database (Accession Number: SRX18100807—18100809). Metagenome raw data have been deposited in NCBI Sequence Short Read Archive (SRA) database (Accession Number: SRX 18100810—18100812).

## ACKNOWLEDGEMENTS

This work was supported by the National Natural Science Foundation of China (91951202) and the National Key R&D Program of China (2018YFA0901200).

## SUPPLEMENTAL MATERIAL

Additional supporting information may be found in the online version of this article at the publisher’s web site.

## Supplementary Methods

### Bacterial community analysis

Microbial community genomic DNA was extracted from sediments samples by using the E.Z.N.A® soil DNA Kit (Omega Biotek) according to the manufacturer’s instructions. The hypervariable regions V3-V4 of bacterial 16S rRNA genes were amplified with the primer pairs <u>338F (5’-ACTCCTACGGGAGGCAGCAG-3’) and 806R (5’-GGACTACHVGGGTWTCTAAT-3’).</u> Purified amplicons were pooled in equimolar and paired-end sequenced on an Illumina MiSeq PE300 platform/NovaSeq PE250 platform (Illumina, San Diego, USA) according to the standard protocols by Majorbio Bio-Pharm Technology Co. Ltd. (Shanghai, China). The raw reads were deposited into the NCBI Sequence Read Archive (SRA) database (Accession Number: SRX18100807-8100809). The raw 16S rRNA gene sequencing reads were demultiplexed, quality-filtered by fastp (version 0.20.0) and merged by FLASH version 1.2.7 with the following criteria: (i) the 300 bp reads were truncated at any site receiving an average quality score of <20 over a 50 bp sliding window, and the truncated reads shorter than 50 bp were discarded, reads containing ambiguous characters were also discarded; (ii) only overlapping sequences longer than 10 bp were assembled according to their overlapped sequence. The maximum mismatch ratio of overlap region is 0.2. Reads that could not be assembled were discarded; (iii) Samples were distinguished according to the barcode and primers, and the sequence direction was adjusted, exact barcode matching, 2 nucleotide mismatch in primer matching.

Operational taxonomic units (OTUs) with 97% similarity cutoff were clustered using UPARSE version 7.1, and chimeric sequences were identified and removed. The taxonomy of each OTU representative sequence was analyzed by RDP Classifier version 2.2 against the 16S rRNA database using confidence threshold of 0.7.

### Metagenomic sequencing and analysis

Microbial community genomic DNA extract was fragmented to an average size of about 400 bp using Covaris M220 (Gene Company Limited, China) for paired-end library construction. Paired-end library was constructed using NEXTflex™ Rapid DNA-Seq (Bioo Scientific, Austin, TX, USA). Adapters containing the full complement of sequencing primer hybridization sites were ligated to the blunt-end of fragments. Paired-end sequencing was performed on Illumina NovaSeq/Hiseq Xten (Illumina Inc., San Diego, CA, USA) at Majorbio Bio-Pharm Technology Co., Ltd. (Shanghai, China) using NovaSeq Reagent Kits/HiSeq X Reagent Kits according to the manufacturer’s instructions (www.illumina.com). Sequence data associated with this project have been deposited in the NCBI Short Read Archive database (Accession Number: SRX 18100810—18100812).

The raw reads from metagenome sequencing were used to generate clean reads by removing adaptor sequences, trimming and removing low-quality reads (reads with N bases, a minimum length threshold of 50bp and a minimum quality threshold of 20) using the fastp (version 0.20.0) on the free online platform of Majorbio Cloud Platform. These high-quality reads were then assembled to contigs using MEGAHIT (version 1.1.2), which makes use of succinct de Bruijn graphs. Contigs with the length being or over 300 bp were selected as the final assembling result.

Open reading frames (ORFs) in contigs were identified using MetaGene (http://metagene.cb.k.u-tokyo.ac.jp/). The predicted ORFs with length being or over 100 bp were retrieved and translated into amino acid sequences using the NCBI translation table SG1. A non-redundant gene catalog was constructed using CD-HIT (version 4.6.1) with 90% sequence identity and 90% coverage. Reads after quality control were mapped to the non-redundant gene catalog with 95% identity using SOAP aligner (version 2.21), and gene abundance in each sample were evaluated. The KEGG annotation was conducted using Diamond (version 0.8.35) against the Kyoto Encyclopedia of Genes and Genomes database (version 94.2) with an e-value cutoff of 1e^−5^.

### Bioinformatics analysis

1904 Rhodobacterales genomes were downloaded from the NCBI database (update to 6 June 2022). The query sequences of SQR, FccB and PDO were from on our previous work. The SQR, FccB, PDO candidate genes in Rhodobacterales genomes were obtained by searching the database with the standalone BLASTP algorithm, using conventional criteria (E value of ≤1e-5, coverage of≥45%, and identity of≥25%). The candidates were analyzed by using ClustalW for alignment and IQ Tree software were used for phylogenetic analysis. The Interactive Tree Of Life was used for the display, manipulation and annotation of phylogenetic trees, and the candidates in the same clade as the seeds were selected. The published flavocytochrome c sulfide dehydrogenase (FCSD) sequences were used as the outgroup of SQR, the published SQR (sulfide:quinone oxidoreductase) sequences were used as the outgroup of FccB, and the GloB (GloB hydroxyacylglutathione hydrolase) sequences were used as the outgroup of PDO.

---

**Figure S1. The main species of sulfane sulfur in sediments is S_8_.** (A) The peak position of S_8_ standard was detected by liquid chromatography. (B) S_8_ increased gradually from the surface layer to the deep layer of sediments.

**Figure S2. The phylogenetic tree of SQRs in Rhodobacterales.** The 344 Rhodobacterales SQR sequences were used for phylogenetic tree construction with reported seed sequences. These sequences were analyzed by using ClustalW, and the tree was built by using IQ Tree. Reference proteins are listed below, some including the organism origin and accession number: BsFCSD, *B. sp*. (ZP_02001369.1); ClFCSD, *C. limicola* (AAL68892.1); RpFCSD, *R. palustris* (YP_001990584.1); AaFCSD, *A. aeolicus* (NP_213158.1). Type I SQRs: RcSqrI, *R. capsulatus* (CAA66112.1); TdSqrI, *T. denitrificans* (AAM52227.1). Type II SQRs: BAO83356.1; AGH39126.1; ABP72874.1; DmSqrII, *D. melanogaster* (NP_647877.1); AmSqrII, *A. marina* (ABV22505.1); MmSqrII, M. musculus (EDL28115.1); SpSqrII, S. pombe (NP_596067.1); PaSqrII, *P. aeruginosa* (NP_251035.1); RsSqrII, *R. solanacearum* (NP_519663.1); ADP99495.1; BAT20392.1; ABI61765.1; ABR69800.1; ALO47562.1; CAM86818.1. Type III SQRs: SsSqrIII, *S. solfataricus* (NP_343961.1); PaSqrIII, *P. aerophilum* (NP_560139.1); AfSqrIII, *A. fulgidus* (NP_069393.1); ABB45153.1; AAU49300.1. Type IV SQRs: SdSqrIV, *S. denitrificans* (YP_393133.1); AvSqrIV, *A. vinosum* (ZP_04773162.1); AEE14738.1; BAF69440.1. Type V SQRs: TvSqrV, *T. volcanium* (NP_111725.1); SsSqrV, *S. solfataricus* (NP_343636.1); StSqrV, *S. tokodaii* (NP_378484.1); TaSqrV, *T. acidophilum* (NP_394588.1). Type VI SQRs: BAP55076.1; AEK59248.1; CCQ75006.1; ADG30032.1; ABK42824.1; TcSqrVI, *T. crunogena* (ABB41976.1); RpSqrVI, *R. palustris* (YP_484673.1). Details of the enzymes and stains can be found in the end of this file.

**Figure S3. The phylogenetic tree of FccBs in *Rhodobacterales*.** The 284 Rhodobacterales FccB sequences were used for phylogenetic tree construction with reported seed sequences. These sequences were analyzed by using ClustalW, and the tree was built by using IQ Tree. Reference proteins are listed below, some including the organism origin and accession number: Type I SQRs: AaSqrI, *A. aeolicus* (NP_214500.1); AhSqrI, *A. halophytica* (AAF72963.1); OlSqrI, *O. limnetica* (AAF72962.1); NsSqrI, *N. sp*. (NP_488552.1); SsSqrI, *S. sp*. (NP_942192.1). Type II SQRs: SpSqrII, *S. pombe* (NP_596067.1); DmSqrII, *D. melanogaster* (NP_647877.1); AmSqrII, *A. marina* (ABV22505.1); MmSqrII, *M. musculus* (EDL28115.1). Type III SQRs: AfSqrIII, *A. fulgidus* (NP_069393.1); MmSqrIII, M. magnetotacticum (ZP_00055086.1); CtSqrIII, *C. tepidum* (NP_661917.1); SsSqrIII, *S. solfataricus* (NP_343961.1); PaSqrIII, *P. aerophilum* (NP_560139.1). Type IV SQRs: SdSqrIV, *S*. *denitrificans* (YP_393133.1); AvSqrIV, *A. vinosum* (ZP_04773162.1); CtSqrIV, *C*. *tepidum* (NP_661023.1). Type V SQRs: TiSqrV, *T. intermedia* (ZP_05500015.1); TvSqrV, *T. volcanium* (NP_111725.1); SsSqrV, *S. solfataricus* (NP_343636.1); StSqrV, *S. tokodaii* (NP_378484.1); CtSqrV, *C. tepidum* (NP_661769.1). Type VI SQRs: TcSqrVI, *T. crunogena* (ABB41976.1); RpSqrVI, *R. palustris* (YP_484673.1); AaSqrVI, *A. aeolicus* (NP_213539.1); ClSqrVI, *C. luteolum* (YP_375032.1); CtSqrVI, *C. tepidum* (NP_661978.1). Subgroup 1 FCCBs: ACL71778.1; CCB65420.1; AKR54467.1; YP_003693897.1; BAR59862.1; YP 001990584.1; ACF00109.1; AMD01363.1; ABE41464.1; CAA55826.2. Subgroup 2 FCCBs: ADD68445.1; ACT47580.1; FCCB YP_286337.1. Subgroup 3 FCCBs: AMO39151.1; ADH65023.1; BAF69150.1. Details of the enzymes and stains can be found in the end of this file.

**Figure S4. The phylogenetic tree of PDOs in *Rhodobacterales*.** The 836 Rhodobacterales PDO sequences were used for phylogenetic tree construction with reported seed sequences. These sequences were analyzed by using ClustalW, and the tree was built by using IQ Tree. Reference proteins are listed below, some including the organism origin and accession number: PaGloB2, *P. aeruginosa* (NP_249523.1); PpGloB2, *P. putida* (ABQ76961.1); EcGloB2, *E. coli* (NP_415447.1); YcbL, *S. enterica* (CAD05397.1); EcGloB1, *E. coli* (NP_414748.1); HiGloB1, *H. influenzae* (ADO96205.1); AtGloB1, *A. fabrum* (NP_356997.2); aGLX2-5, *A. thaliana* (NP_850166.1); ytGLO_2_, *S. cerevisiae* (CAA71335.1); hGLX2, *H. sapiens* (CAA62483.1); BcII, B. cereus (AAA22276.1). Type I PDOs: MxPdoI (YP_633997.1), HsPdoI (NP_055112.2), BvPdoI (ZP_00420127.1), AtPdoI (NP_974018.3); SaPdoI (YP_003957083.1); AFS48613.1. Type II PDOs: ACL93948.1; SmPdoII (NP_435818.1), AfaPdoII (AAK89929.1), XfPdoII (NP_298058.1), AMB74311.1; AJE45940.1; ABP72383.1; CpPdoII (YP_297791.1), PaPdoII (NP_251605.1), BxPdoII (YP_554628.1); CAP57776.1. Type III PDOs: SaPdoIII, *S.aureus* (WP_000465474.1); BAS29225.1; ABE45829.1; ADG06957.1; ACZ43611.1; BAL99983.1; AMM91879.1; AKG72915.1; AEA26511.1; ACY97605.1; ADI04923.1. Details of the enzymes and stains can be found in the end of this file.

**Table S1.** Strains and plasmids used in this study.

**Table S2.** Primers used for gene deletion.

**Table S3.** Primers used for real-time PCR.

**Table S4.** Sulfur oxidation genes in 1904 Rhodobacterales.

